# The Perfect Storm: Gene Tree Estimation Error, Incomplete Lineage Sorting, and Ancient Gene Flow Explain the Most Recalcitrant Ancient Angiosperm Clade, Malpighiales

**DOI:** 10.1101/2020.05.26.112318

**Authors:** Liming Cai, Zhenxiang Xi, Emily Moriarty Lemmon, Alan R. Lemmon, Austin Mast, Christopher E. Buddenhagen, Liang Liu, Charles C. Davis

## Abstract

The genomic revolution offers renewed hope of resolving rapid radiations in the Tree of Life. The development of the multispecies coalescent (MSC) model and improved gene tree estimation methods can better accommodate gene tree heterogeneity caused by incomplete lineage sorting (ILS) and gene tree estimation error stemming from the short internal branches. However, the relative influence of these factors in species tree inference is not well understood. Using anchored hybrid enrichment, we generated a data set including 423 single-copy loci from 64 taxa representing 39 families to infer the species tree of the flowering plant order Malpighiales. This order alone includes nine of the top ten most unstable nodes in angiosperms, and the recalcitrant relationships along the backbone of the order have been hypothesized to arise from the rapid radiation during the Cretaceous. Here, we show that coalescent-based methods do not resolve the backbone of Malpighiales and concatenation methods yield inconsistent estimations, providing evidence that gene tree heterogeneity is high in this clade. Despite high levels of ILS and gene tree estimation error, our simulations demonstrate that these two factors alone are insufficient to explain the lack of resolution in this order. To explore this further, we examined triplet frequencies among empirical gene trees and discovered some of them deviated significantly from those attributed to ILS and estimation error, suggesting gene flow as an additional and previously unappreciated phenomenon promoting gene tree variation in Malpighiales. Finally, we applied a novel method to quantify the relative contribution of these three primary sources of gene tree heterogeneity and demonstrated that ILS, gene tree estimation error, and gene flow contributed to 15%, 52%, and 32% of the variation, respectively. Together, our results suggest that a perfect storm of factors likely influence this lack of resolution, and further indicate that recalcitrant phylogenetic relationships like the backbone of Malpighiales may be better represented as phylogenetic networks. Thus, reducing such groups solely to existing models that adhere strictly to bifurcating trees greatly oversimplifies reality, and obscures our ability to more clearly discern the process of evolution.

## Introduction

One of the most difficult challenges in systematics is reconstructing evolutionary history during periods of rapid radiation. During such intervals, few DNA substitutions accrue, rendering little information for phylogenetic inference. The potentially large population sizes and close evolutionary relationships create opportunities for widespread incomplete lineage sorting (ILS) and gene flow, leading to excessive gene tree-species tree conflict. The tremendous growth of genome-scale data sets, however, has greatly improved researchers’ ability to investigate rapid radiations by providing hundreds to thousands of unlinked loci. Commonly applied approaches include not only whole genome sequencing but also RNA-Seq, RAD-Seq, and anchored hybrid enrichment, which in general are cost effective and efficient across broad taxonomic groups and yield data sets with dense locus and taxon sampling (Lemmon and Lemmon 2013). These approaches are promising and have been variously applied to successfully resolve a number of recalcitrant clades across the Tree of Life, including in birds (Prum et al. 2015), mammals (Song et al. 2012), fish (Wagner et al. 2013), and plants (Wickett et al. 2014).

Despite their promise, however, these enormous data sets also introduce new methodological challenges and complexities. In particular, phylogenomic data sets may yield strongly supported, yet conflicting or artifactual, results depending on the method of inference or genomic regions sampled (Song et al. 2012; Jarvis et al. 2014; Xi et al. 2014; Reddy et al. 2017; Shen et al. 2017). During rapid radiations, ILS can lead to extreme conditions where the most probable gene tree differs from the topology of the true species tree, which is referred to as the “anomaly zone” (Degnan and Rosenberg 2006; Rosenberg and Tao 2008). Such pervasive genealogical discordance, in particular, can result in biased species tree inference when applying concatenation methods, and produce inconsistent and conflicting results with strong confidence (Song et al. 2012; Xi et al. 2014). The multispecies coalescent (MSC) model, which explicitly accommodates gene tree heterogeneity caused by ILS, in contrast, has been demonstrated to be more reliable under these circumstances. Most recently, a class of “two-step” summary coalescent methods has been the focus of substantial development and application (Nakhleh 2013). They are demonstrated to be statistically consistent under the MSC model and can work efficiently with genome-scale data (Liu et al. 2009; Liu et al. 2010; Chifman and Kubatko 2014; Mirarab et al. 2014c). Their application has been successful in resolving mammalian, avian, and seed plant relationships in cases where concatenation methods have been demonstrated to be inconsistent (Song et al. 2012; Xi et al. 2013; Reddy et al. 2017).

In addition to ILS, gene tree estimation error has also been a major focus of work to improve the accuracy of phylogenomic inference. This is especially relevant for summary coalescent methods, which assume the input gene trees to be essentially error-free, e.g., Lanier et al. 2014; Mirarab et al. 2014c; Roch and Warnow 2015; Xu and Yang 2016; Blom et al. 2017. Rapid radiations are particularly challenging in this regard. Here, short internal branches may yield error-prone gene tree estimation when phylogenetically informative characters are minimal (Xi et al. 2015). This may be further complicated if such radiations are ancient and followed by long descendent branches, which may exacerbate long-branch attraction artifacts (Whitfield and Kjer 2008). Though benchmark studies have demonstrated the consistency of summary coalescent methods when substantial amounts of such non-phylogenetic signal are included (Philippe et al. 2011; Roch and Warnow 2015; Xi et al. 2015; Hahn and Nakhleh 2016), accurate gene tree inference remains of crucial importance for reliable species tree estimation (Shen et al. 2017). A number of methods have been developed to mitigate gene tree estimation error, including improving taxon sampling, applying appropriate models of nucleotide evolution, reducing missing data, subsampling informative genes, and locus binning (Zwickl and Hillis 2002; Lemmon et al. 2009; Salichos and Rokas 2013; Cox et al. 2014; Mirarab et al. 2014a; Hosner et al. 2015).

Beyond ILS and gene tree estimation error, gene flow between non-sister species can similarly result in gene tree–species tree conflict and lead to incorrect species tree estimation. Unlike the MSC model, gene flow from a non-sister species leads to an overrepresentation of the parental allele in the descendants and therefore the frequencies of the two minor topologies are asymmetrical (Durand et al. 2011). A number of species network inference methods have been developed to detect and infer gene flow based on such expectation. They either use counts of the shared derived alleles, such as the classic D-statistic test (Green et al. 2010; Durand et al. 2011), or the gene tree topology as input (e.g., Huson et al. 2005; Meng and Kubatko 2009; Yu et al. 2011; Solís-Lemus et al. 2017). The latter methods are often based on *a priori* evolutionary models and have been increasingly applied to empirical data sets.

During periods of rapid radiation, all of the above phenomena—ILS, introgression, and gene tree estimation error—may occur simultaneously to obscure phylogenetic signal (Pease et al. 2016), culminating in a perfect storm confounding phylogenomic inference. When a limited number of alternative species tree topologies are involved, these phenomena can be distinguished from each other using methods discussed above (Zwickl et al. 2014; Arcila et al. 2017; Meyer et al. 2017; Beckman et al. 2018; Glémin et al. 2019). However, when the rapid radiation generates a cloud of alternative tree topologies, all of which are weakly supported, such model-based methods become less practical because priors necessary to test hypothesis of introgression are difficult to determine accurately. Additional challenges arise from the excessive computational resources required to apply such network inference methods to data sets involving hundreds of species. Moreover, following the identification of ILS, introgression, and gene tree estimation error, a more quantitative assessment characterizing their relative contribution to overall gene tree variation has not been addressed in any empirical system to our knowledge.

Using anchored hybrid enrichment (Lemmon et al. 2012), we generated a large phylogenomic data set including 423 single-copy nuclear loci with 64 taxa to infer relationships of the flowering plant clade Malpighiales. The order Malpighiales comprise ca 7.8% of eudicot diversity (Magallon et al. 1999) and include more than 16,000 species in ~36 families (Stevens and Davis 2001). Species in Malpighiales encompass astonishing morphological and ecological diversity ranging from epiphytes (Clusiaceae), submerged aquatics (Podostemaceae), to emergent rainforest canopy species (Callophyllaceae). The order also includes numerous economically important crops with sequenced genomes, e.g., rubber (*Hevea*), cassava (*Manihot*), flax (*Linum*), and aspen (*Populus*). Despite their ecological and economic importance, the evolutionary history of Malpighiales remains poorly understood. While analyzing chloroplast genome sequences has greatly improved the resolution of this clade, relationships among its major subclades remain uncertain (Xi et al. 2012), and analyses using nuclear genes lack resolution along the spine of the clade (Davis et al. 2005; Wurdack and Davis 2009). According to Smith et al. (2013), this region of the Malpighiales phylogeny has been implicated in nine of the top ten most unstable nodes across all angiosperms, including Pandaceae, Euphorbiaceae, Linaceae, the most recent common ancestor (MRCA) of Salicaceae and Lacistemataceae, the MRCA of Malpighiaceae and Elatinaceae, as well as the MRCA of putranjivoids, phyllanthoids, chrysobalanoids, and rhizophoroids *sensu* Xi et al. (2012). In short, Malpighiales have been coined one of the “thorniest nodes” in the angiosperm tree of life (Soltis et al. 2005). A longs-tanding hypothesis for this lack of resolution has been attributed to the clade’s rapid radiation during the Albian and Cenomanian (112–94 million years ago [Ma]; Davis et al. 2005; Wurdack and Davis 2009; Xi et al. 2012). This radiation has produced a phylogeny characterized by extremely short internal branches along the backbone of the phylogeny, followed by long branches subtending most crown group families. This is particularly problematic because, as we summarize above, short internal branches represent species tree anomaly zones where ILS may be pervasive and gene tree estimation error is high (Liu et al. 2015; Roch and Warnow 2015; Edwards et al. 2016). Incongruent phylogenetic signals between organelle and nuclear genes also support introgression associated with the origin of this order (Sun et al. 2015).

The development of next-generation sequencing, the MSC model that accommodate ILS, and best practices to reduce gene tree estimation error offers a unique opportunity to re-examine Malpighiales in the context of resolving rapid radiations. Here, we apply both concatenation and coalescent-based methods for phylogenomic analyses and evaluate the relationships and consistency of nodal resolution under a variety of conditions. We also apply simulations to explore the impact of ILS and gene tree estimation error based on the empirical parameters of our inferred species tree. We further apply a triplet analysis to detect gene flow and identify hotspots of reticulate evolution in the species tree. And finally, we develop a novel method to quantitatively assess the contribution of three primary sources of gene tree variation in Malpighiales—ILS, gene tree estimation error, and gene flow.

## Materials And Methods

### Taxon Sampling

We sampled a total of 56 species in the order Malpighiales, representing 39 families and all major clades *sensu* Wurdack and Davis (2009) and Xi et al. (2012) (Table S1). Species were sampled to represent the breadth of Malpighiales diversity. Four species from the order Celastrales and two species from the order Oxalidales were sampled as closely related outgroups (Chase et al. 2016). Two species from the order Vitales were also included as more distantly related outgroups (Chase et al. 2016, Table S1).

### Library Preparation, Enrichment, and Locus Assembly

Data were collected at the Center for Anchored Phylogenomics at Florida State University (http://www.anchoredphylogeny.com) using the anchored hybrid enrichment method (Lemmon et al. 2012; Buddenhagen et al. 2016). This method targets universally conserved single-copy regions of the genome that typically span 250 to 800 base pairs (bp), thus mitigating the confounding effect of paralogy in gene tree estimates. Briefly, total genomic DNA was sonicated to a fragment size of 300–800 bp using a Covaris E220 Focused-ultrasonicator. Library preparation and indexing was performed following the protocol in Hamilton et al. (2016). A size-selection step was also applied after blunt-end repair using SPRI select beads (Beckman-Coulter Inc). Indexed samples were then pooled and enriched using the Angiosperm v1 kit (Agilent Technologies Custom SureSelect XT kit ELID 623181; Buddenhagen et al. 2016). The resulting libraries were sequenced on an Illumina HiSeq 2500 System using the PE150 protocol.

Quality-filtered sequencing reads were processed following the methods described in Hamilton et al. (2016) to generate locus assemblies. Briefly, paired reads were merged prior to assembly following Rokyta et al. (2012). Reads were then mapped to the probe region sequences of the following reference genomes: *Arabidopsis thaliana* (Malvales, Arabidopsis Genome Initiative 2000), *Populus trichocarpa* (Malpighiales, Tuskan et al. 2006), and *Billbergia nutans* (Poales, Buddenhagen et al. 2016). Finally, the assemblies were extended into the flanking regions. Consensus sequences were generated from assembly clusters with the most common base being called when polymorphisms could be explained as sequencing error.

### Orthology Assignment

Orthologous sequences were determined following Prum et al. (2015) and Hamilton et al. (2016). The assembled sequences were grouped by locus and a pairwise distance was calculated as the percent of shared 20-mers. Sequences were subsequently clustered based on this distance matrix using the neighbor-joining algorithm (Saitou and Nei 1987). When more than one cluster was detected for a target region, each cluster was treated as a different locus in subsequent analyses. Clusters including less than 50% of the species were discarded.

### Sequence Alignment, Masking, and Site-subsampling

Each locus was first aligned using MAFFT v7.023b (Katoh and Standley 2013) with “--genafpair --maxiterate 1000” flags imposed. Alignments were end trimmed and internally masked to remove misassembled or misaligned regions (Buddenhagen et al. 2016). Firstly, conserved sites were identified in each alignment where >40% of the nucleotides at that site were identical across species. For end trimming, sequences for each gene accession were scanned from both ends towards the center until more than fourteen nucleotides in a sliding window of 20 bp matched the conserved sites. Once the start and end of each sequence was established, the internal masking then required that >50% of the nucleotides in a sliding window of 30 bp matched the conserved sites. Regions that did not meet this criterion were masked. Finally, we removed any gene sequence in the alignment with >50% ambiguous nucleotide composition. We also required all locus alignments to contain *Leea guineense* (Vitales) for rooting purposes.

To further explore the phylogenetic utility of the flanking regions of hybrid enrichment data, we applied three increasingly stringent site-subsampling strategies using trimAl v1.2 (Capella-Gutiérrez et al. 2009) following our masking steps described above. To construct our “low-stringency data set”, we set the gap threshold to be 0.8 (−gt 0.8) in trimAl to remove sites containing >20% indels or missing data for each alignment. This data set includes the highest percentage of flanking regions and resulted in the longest alignments. We then applied a site composition heterogeneity filter to this “low-stringency data set” to create our “medium-” and “high-stringency data set” by setting the minimum site similarity score to be 0.0002 and 0.001 (e.g., −st 0.001), respectively. This has the effect of removing especially rapidly evolving sites within flanking regions for which we expect higher composition heterogeneity. The resulting “medium-” and “high-stringency data set” thus include lower percentage of flanking regions.

### Gene Tree Estimation

To infer individual gene trees for coalescent-based analyses, we applied maximum likelihood (ML) as well as Bayesian Inference (BI). To estimate ML trees, we used RAxML v8.1.5 (Stamatakis 2014) under the GTR+Γ model with 20 random starting points. We chose the GTR+Γ model because it accommodates rate heterogeneity among sites, while the other available GTR model in RAxML, the GTRCAT model, is less appropriate due to our small taxon sampling size (Stamatakis 2014). Statistical confidence of each gene tree was assessed by performing 100 bootstrap (BP) replicates. We additionally inferred the Bayesian posterior distribution of gene trees using MrBayes v3.2.1 (Ronquist and Huelsenbeck 2003). We only applied BI to the low-stringency data set due to computational cost and this data set yielded the best resolved gene trees (see Results below). We applied the GTR+Γ model with two independent runs for each gene. Each run included four chains, with the heated chain at temperature 0.20 and swapping attempts every 10 generations. Initially, four million generations were used with 25% burn-in period, sampled every 1,000 generations. Runs that failed to reach the targeted standard deviation of split frequencies ≤0.02 were rerun with the same settings but with 10 million generations, sampled every 5,000 generations until attaining a standard deviation of split frequencies ≤0.02. We randomly sampled 100 trees in the posterior distribution of inferred gene trees to conduct bootstrap replication in the coalescent analyses (Table S2). Trees sampled from the posterior distribution are more similar to the optimum Bayesian tree than those sampled from the non-parametric bootstrapping. Therefore, we also expect higher support values in the species tree.

### Species Tree Inference Using Concatenation and Coalescent-based Methods

Our trimmed gene matrices were concatenated and analyzed using both RAxML and ExaML v3.0.18 (Kozlov et al. 2015). In our RAxML analyses, the species trees were inferred under the GTR+Γ model with 100 rapid bootstrapping followed by a thorough search for the ML tree. In ExaML analyses, species trees were inferred under the GTR+Γ model with 20 random starting points. We then conducted 100 bootstrap replicates to evaluate nodal support. Partitions for both analyses were selected by PartitionFinder v2.1.1 based on AICc (Akaike Information Criterion) criteria using the heuristic search algorithm “rcluster” (Lanfear et al. 2012). We also conducted BI for species tree estimation as implemented in PhyloBayes (Lartillot et al. 2013). For BI analysis, we applied the CAT-GTR model, which accounts for across-site compositional heterogeneity using an infinite mixture model (Lartillot and Philippe 2004). Two independent Markov chain Monte Carlo (MCMC) analyses were conducted for each concatenated nucleotide matrix. Convergence and stationarity from both MCMC analyses were determined using bpcomp and tracecomp from PhyloBayes. We ran each MCMC analysis until the largest discrepancy observed across all bipartitions was smaller than 0.1 and the minimum effective sampling size exceeded 200 for all parameters in each chain.

To infer our species tree using coalescent-based models, we obtained ML gene trees and BI consensus trees for each locus. MP-EST (Liu et al. 2010) and ASTRAL-II (Mirarab and Warnow 2015) were subsequently used to perform species tree inference using optimally estimated gene trees. Statistical confidence at each node was evaluated by performing the same species tree inference analysis on 100 ML bootstrap gene trees or trees sampled from our Bayesian posterior distributions. The resulting 100 species trees estimated from bootstrapped samples were summarized onto the species tree inferred from ML gene trees using the option “-f z” in RAxML.

### Simulation of gene alignments with realistic parameters of ILS and gene tree estimation error

To investigate the impact of ILS and gene tree estimation error on the accuracy of species tree inference we simulated sequences assuming a known species tree. Here, the tree topology estimated by MP-EST with the low-stringency data set (analysis No. 15 in Table S2) was invoked as the known species tree. We chose this best-supported MP-EST topology because the branch lengths are estimated in coalescent unit, which is an essential parameter for ILS simulation. We thus applied this species tree to all of the downstream simulation-based analyses, including the triplet test for MSC model fitness and relative importance analysis.

To simulate conditions of high and low levels of ILS, we modified the key population mutation parameter “theta” when generating gene trees under the coalescent model using the function “sim.coaltree.sp.mu” in the R package Phybase (Liu and Yu 2010). Theta was set to be 0.01 and 0.1 to reflect low and high ILS, respectively. The range of theta was determined based on our empirical data sets by following two steps. First, we inferred the branch lengths of the species tree in mutation units in RAxML using the fixed topology of the MP-EST species tree and the concatenated low-stringency data set. Second, theta for each branch was calculated by dividing the branch lengths estimated from RAxML (mutation units) by that estimated from MP-EST (coalescent units). The other input for Phybase, the ultrametric species tree, was generated from this RAxML phylogeny using the function “chronos” in the R package ape (Paradis et al. 2004). In addition, we set the relative mutation rates to follow a Dirichlet distribution with alpha equal to 5.0. This alpha reflected the large variance in gene mutation rates. Finally, 1,500 non-ultrametric gene trees were simulated separately for each theta.

From these simulated gene trees, DNA alignments of different lengths were subsequently generated to reflect various levels of gene tree estimation error since alignment length is easy to manipulate and shorter alignments correspond to higher error rates (Mirarab et al. 2014b). We used bppsuite (Guéguen et al. 2013) to simulate alignments under the GTR+Γ model. Parameters of the model, including the substitution matrix, base frequency, and the gamma rate distribution were extracted from the RAxML phylogeny above inferred from the low-stringency data set. For each gene tree we generated alignments of 300, 400, 500, 1,000, and 1,500 bp.

As a result, fifty data sets were generated by including 100, 200, 500, 1,000, and 1,500 simulated loci of five length categories and two theta categories (Table S3, Fig. 1). Species trees were inferred using the concatenation and coalescent methods as described above under these varying levels of ILS and gene tree estimation error. Finally, we quantified gene tree–species tree discordance and species tree error by measuring the RF distance between an estimated gene tree or species tree to the true species tree.

**Figure 1.**
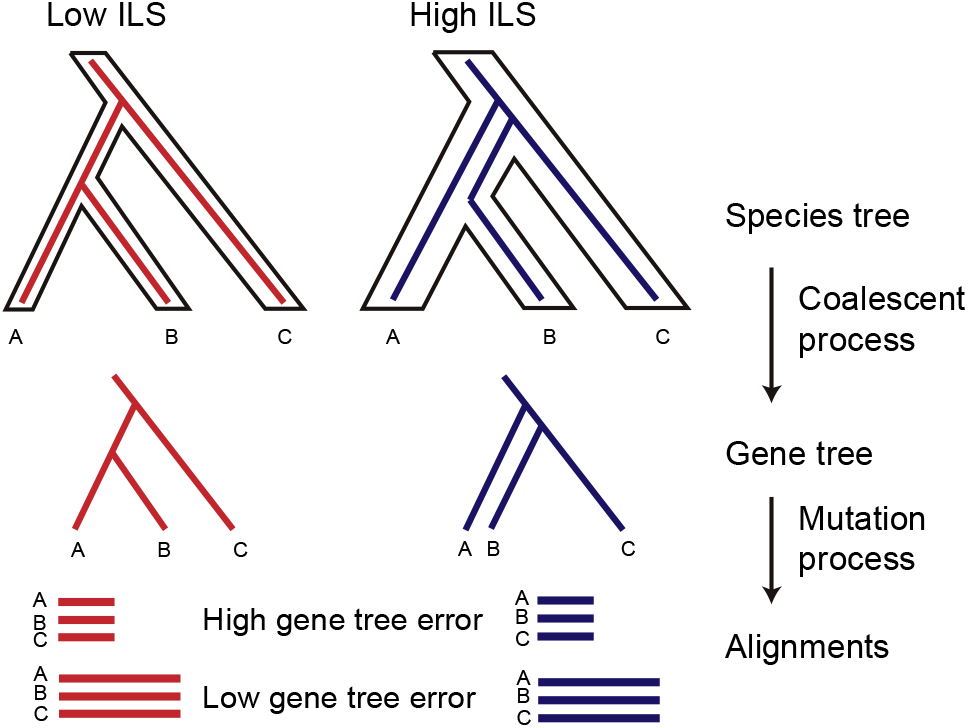
Simulation of ILS and gene tree estimation error. ILS was simulated though the coalescent process by setting low (0.01) and high (0.1) theta values. DNA alignments were subsequently generated through the mutation process based on simulated gene trees. Five alignments were generated for each gene tree with lengths of 300, 400, 500, 1000, and 1500 bp (only two are shown in the graph). Shorter alignment lengths increase in gene tree estimation error.

In order to assess the sensitivity of our simulation results to the choice of input species tree and theta values, we additionally examined gene tree–species tree discordance among bootstrapped samples. We simulated 1,500 gene trees for each of the 100 MP-EST bootstrapped species trees. Gene trees were simulated directly from each species tree using the “sim.coal.mpest” function in the R package Phybase (Liu and Yu 2010). This method does not require *a priori* theta parameters as was imposed in our simulation above and so alleviates concerns of applying erroneous theta values. We subsequently quantified the gene tree–species tree discordance for each bootstrap replicate as described above. We did not use these gene trees to simulate alignments because these gene trees are ultrametric (Liu and Yu 2010) and thus not suitable for such purpose.

### A Test of the MSC Model Using Triplet Frequencies

To determine the fit of the MSC model to our empirical data we additionally examined the triplet frequency for all 423 ML genes trees inferred from our low-stringency data set using a custom R script available on Github (http://github.com/lmcai/Coalescent_simulation_and_gene_flow_detection). We used the asymmetrical triplet frequency as evidence for introgression (Fig. 2a). This metric has been widely applied in parsimony, likelihood, and Bayesian based species network inference methods to detect sources of gene flow (Nakhleh 2013). Our method differs from these methods in two aspects: first, the statistical significance of asymmetry in triplet frequency is determined by a null distribution simulated from the empirical data. We took into account variations from ILS and missing data, thus reducing the false positive rate. Second, unlike other model-based species network inference methods, after identifying significantly asymmetrical triplets, we used a novel method to summarize and visualize the distribution of lineages involved in gene flow on a species tree without optimizing the global network (Fig. 2b). As a result, our methods can be easily scaled to genomic data involving hundreds of taxa.

**Figure 2.**
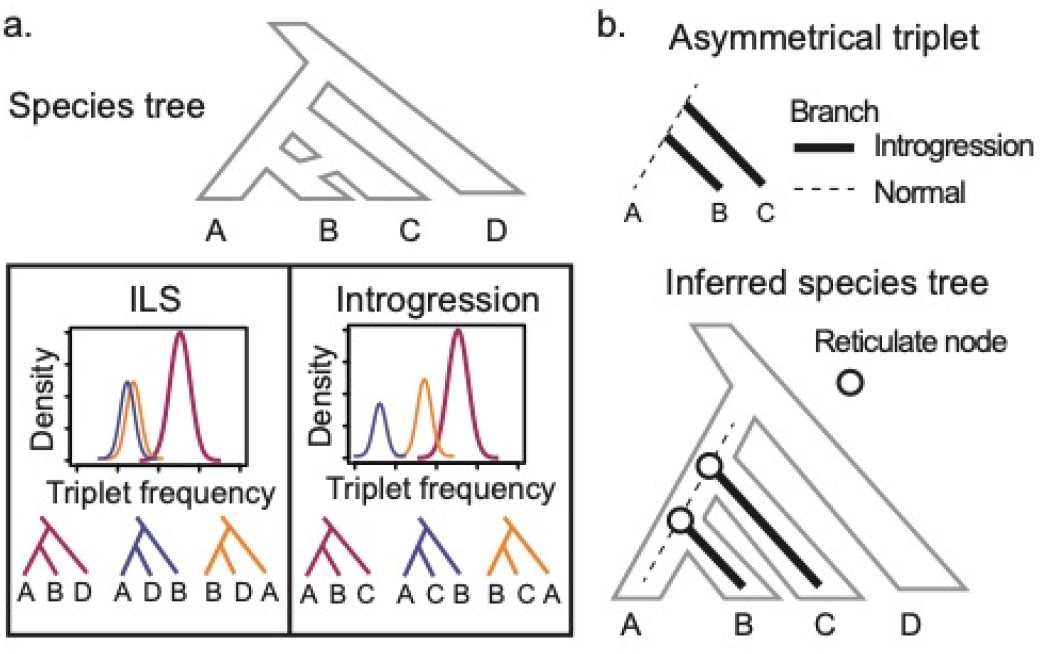
Identification of reticulate evolution using triplet frequency. (a) Theoretical expectations of triplet frequency distribution under the multi-species coalescent (MSC) model with and without introgression. In case of incomplete lineage sorting (ILS), symmetrical distributions of the frequency of two minor topologies are excepted owing to deep coalescence (left). In case of introgression, one of the minor topologies will occur with higher frequency due to gene flow (right). (b) Mapped asymmetrical triplets to species tree to identify reticulate nodes.

In order to identify a triplet with significantly asymmetrical frequencies, we generated a null distribution of triplet frequencies for each triplet using simulated gene trees under the MSC model. For each of the 100 MP-EST BP species trees, we simulated 423 gene trees using the “sim.coal.mpest” function in Phybase. For each set of simulated gene trees, we then generated missing data for each species by pruning that species randomly among all gene trees so that the number of sampled genes of that species was the same as the empirical data. We subsequently counted triplet frequency for these gene trees in each bootstrap replicate. This simulated distribution reflects the variation of triplet frequency arising from ILS, estimation error, sampling error, and missing data. A triplet in the empirical data was identified to be significantly asymmetrical if the difference between the two less frequent triplets exceeded the maximum difference under simulated conditions. Such triplets potentially violate the assumptions of the MSC model, and point towards gene flow especially as an additional factor influencing gene tree heterogeneity, though ancestral population structure (Slatkin and Pollack 2008) and biases in substitution or gene loss can produce asymmetrical triplet as well (see Discussion below).

### Identifying hotspots of reticulate evolution using the Reticulation Index

We developed a relative measurement statistic, the ‘Reticulation Index’, to quantify the intensity of introgression at each node. First, for each asymmetrical triplet, we mapped the two inferred introgression branches to the species tree (Fig. 2b). Second, for each node on the species tree, we counted the number of introgression branches that were mapped to it. These raw counts were then normalized by the total number of triplets associated with that node. The resulting percentage is the Reticulation Index for each node. The R script for calculating the Reticulation Index and visualizing the result on a species tree is available in the above Github repository.

### A Novel Method to Quantify Gene Tree Variation Due to ILS, Gene Tree Estimation Error, and Gene Flow

Untangling the effects of ILS, gene tree estimation error, and gene flow is challenging since they all lead to gene tree–species tree discordance. Here, based on a multiple regression model (Grömping 2006), we assign shares of relative importance to ILS, gene tree estimation error, and gene flow in generating gene tree variation by variance decomposition.

For all 63 internal nodes in our species tree, we separately estimated the level of ILS, gene tree estimation error, and gene flow for each node. ILS is represented by our estimates of theta. Gene flow is represented by the Reticulation Index for each node. To infer the level of gene tree estimation error at each node, we additionally simulated 423 gene alignments of 446 bp (median alignment length in low-stringency data set) from the MP-EST species tree, but each with unique substitution model parameters estimated from the empirical alignments. This simulation and phylogeny inference followed the same strategy of alignment simulation described above (Fig. 1). We subsequently inferred phylogenies for these alignments and summarized them on the species tree to obtain the BP value at each node. Here, the BP values represent the gene tree variation generated by estimation error.

The gene tree variation in the empirical data is obtained by summarizing bootstrap trees from each of the 423 loci in our low-stringency data set onto the species tree. The resulting BP values represented observed gene tree variation at each node. We then inferred the relative contribution of ILS, estimation error, and gene flow in explaining gene tree variation using linear regression methods implemented in the R package relaimpo (Grömping 2006). We used four different methods, “lmg”, “last”, “first”, and “Pratt”, to decompose the relative importance of the three regressors (Lindeman 1980; Pratt 1987). All of these methods are capable of dealing with correlated regressors and “lmg” is the most robust method among them (Grömping 2006). We applied the functions “boot.relimp” and “booteval.relimp” to estimate the relative importance and their confidence interval by bootstrapping 100 times.

### Testing the utility of the triplet-frequency-based method using a genomic data set from yeast

To further validate our introgression detection method using the triplet frequency distribution, we applied it to the benchmark multi-locus yeast data set from Salichos and Rokas (2013). We obtained the 1,070 gene trees and inferred a species tree using MP-EST. We also conducted 100 bootstrap replicates of species tree inference using the bootstrap gene trees. We then applied our triplet method to identify asymmetrical triplets as described above (*A Test of the MSC Model Using Triplet Frequencies*). Finally, all asymmetrical triplets were mapped to the inferred species tree and the Reticulation Index for each node was calculated and visualized as described above (*Identifying hotspots of reticulate evolution using the Reticulation Index*).

## Results

### Hybrid Enrichment

We successfully captured and sequenced 423 of our 491 targeted loci. The resulting data matrix was densely sampled and included only 12% missing data. One hundred and one loci included at least 61 taxa (>95% occupancy) and only four loci had more than 19 missing species (>30%). The locus sampling per taxon varied from 423 (*Leea guineense*) to 278 (*Ouratea sp*. and *Lophopyxis maingayi*, Table S4). After applying site subsampling, the alignment lengths ranged from 190 to 885 bp (median 446 bp) for the low-stringency data set, 157 to 791 bp (median 376 bp) for the medium-stringency data set, and 112 to 751 bp (median 271 bp) for the high-stringency data set (Table S4). In all data sets, the number of parsimony informative sites and the average nodal support was significantly positively correlated with alignment length (*p*-value <1e-5, Fig. S1).

### Flanking Regions Increase Gene Tree and Species Tree Resolution

We observed increasing mean BP support among gene trees as increasingly larger percentages of the flanking region were included. The average gene tree nodal support from our low-stringency data set (42 ML BP) was significantly higher than nodal support estimates for the medium (39 ML BP, *p*-value = 1.2e-89 in paired t-test) and high-stringency data sets (35 ML BP, *p*-value = 1.3e-77, Fig. S2a).

These increases in gene tree resolution also contributed to increased species tree resolution as well as species tree inference congruency. For both concatenation and coalescent analyses, species trees estimated from the low stringent data set with highest amount of flanking regions, always resulted in the highest average BP support (Fig. S2b,c, Table S2) and the lowest pairwise RF distances (Fig. S2d) indicating increased statistical consistency when adding flanking regions.

### Malpighiales Species Tree Resolution

We observed significantly higher average species tree nodal support in concatenation compared to coalescent reconstructions (Table S2, *p*-value = 2.62e-28 in paired *t*-test). However, our results also suggest statistical inconsistency across data sets when applying concatenation (Fig. S3). The higher pairwise weighted Robinson–Foulds distance (WRF) in concatenation indicate more well-supported conflicts among species trees, which further supports mounting evidence that coalescent methods are more consistent when reconstructing species tree relationships involving extensive ILS (i.e., the anomaly zone, Degnan and Rosenberg, 2006, Rosenberg and Tao, 2008). In addition, we did not find locus subsampling based on locus length, number of PI sites, or gene tree quality help increase species tree resolution (see Supplementary Note 1, Table S2).

Our most well resolved species trees estimation inferred with ASTRAL and MP-EST uncovered ten major subclades of Malpighiales (Clade 1 to 10 in Fig. 3). These relationships corresponded to families or closely related clades of families, five of which have previously been identified using plastid genome (Fig. 3, Xi et al. 2012). Five new clades were supported with ≥50 BP, >0.90 PP. Three of these newly identified clades are in conflict (>70 BP) with the plastid phylogeny from Xi et al. (2012) and are discussed more extensively below. Interrelationships among these ten major subclades, however, were not well resolved (<50 BP).

**Figure 3.**
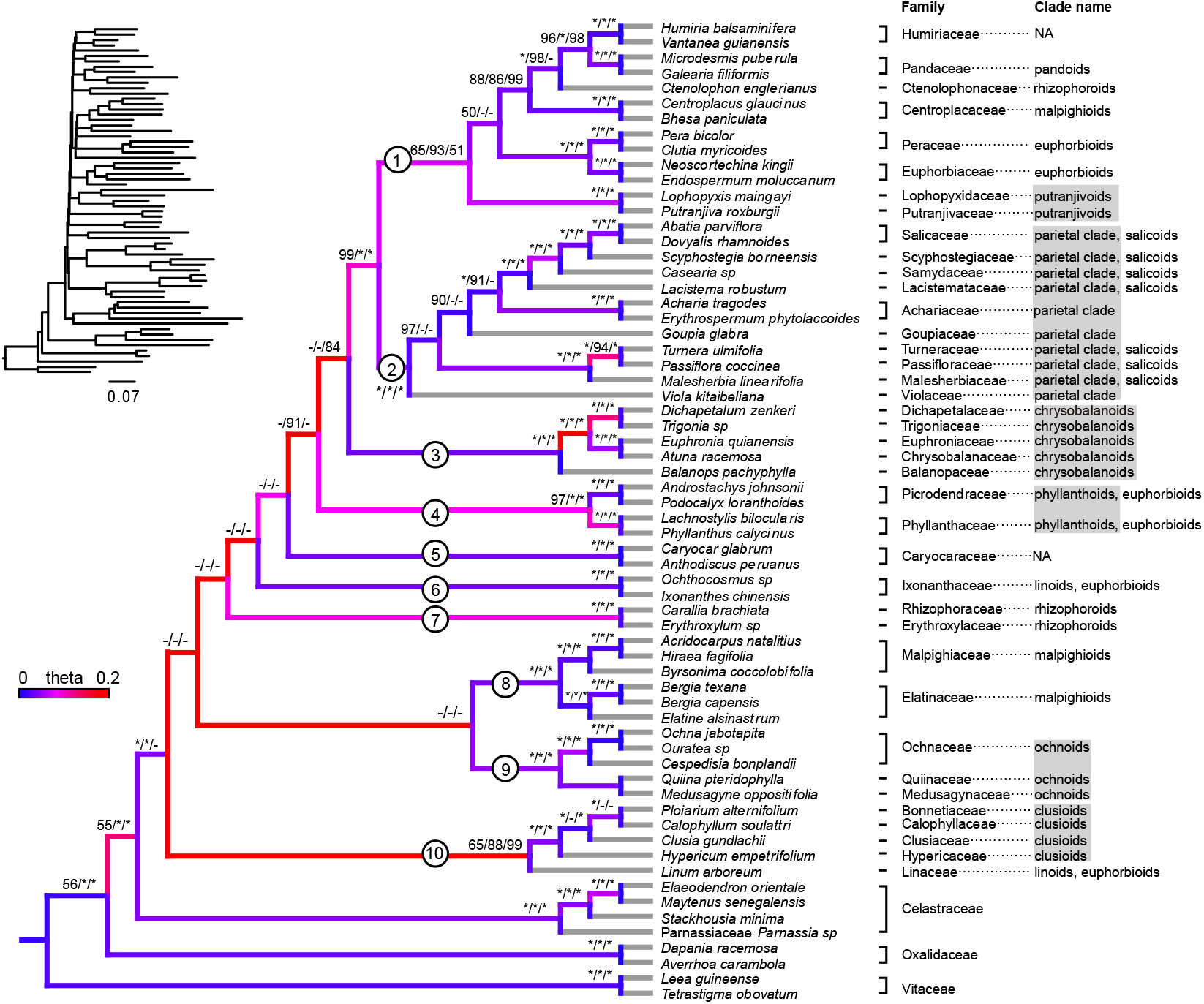
Species phylogeny of Malpighiales derived from MP-EST with complete low-stringency data set (analysis No. 15 in Table S2). Gene trees are estimated using MrBayes. Branches are colored by the inferred population mutation parameter theta. Warmer colors indicate higher theta and thus higher level of ILS. Terminal branches are colored grey due to lack of data to infer theta. BP values from best-resolved MP-EST/ASTRAL/RAxML analyses (analysis No. 15, 17, and 11 in Table S2) are indicated above each branch; an asterisk indicates 100 BP support; a hyphen indicates less than 50 BP. Branch lengths estimated from RAxML by fixing the species tree topology are presented at the upper left corner. The eleven major clades highlighted in the discussion are identified with circled numbers along each relevant branch. The clade affiliation for each family based on the plastid phylogeny (Xi et al. 2012) is indicated on the right. Clades identified by Xi et al. (2012) that are also monophyletic in this study are highlighted using gray shades.

### Simulated Levels of ILS and Gene Tree Estimation Error Reflects Empirical Data

The 5% and 95% quantiles of theta were inferred to be 0.0254 and 0.176, respectively, with a median of 0.0688. High theta was mostly found along the backbone of the species tree, indicating the likelihood of extensive ILS within this region of the tree (Fig. 3). This is likely an overestimation of theta since all topological variations are attributed to coalescent process including the ones originate from mutational variance (Huang and Knowles 2009). We therefore set the theta parameter to be 0.01 and 0.1 in our coalescent gene tree simulation, which reflected the left and right tails of low and high ILS estimated from empirical data.

In our simulation, the average gene tree estimation error was 0.319 for alignments of 300bp, 0.261 for 400bp, 0.221 for 500bp, 0.133 for 1000bp and 0.098 for 1500bp under low ILS and 0.340 for alignments of 300bp, 0.286 for 400bp, 0.241 for 500bp, 0.161 for 1000bp and 0.120 for 1500bp under high ILS. Here, an RF distance of 0 signifies error-free reconstruction versus 1 indicating that none of the true nodes are recovered. Gene tree estimation error was therefore lower in low ILS (*p*-value=6.08e-16 in Student’s *t*-test), but was still significantly higher than that estimated from empirical data (*p*-value= 4.24e-65 in Student’s *t*-test; see Supplementary Note 2; Fig. S4).

### Simulation Yields Consistent and Accurate Species Tree Estimation

In our empirical analyses, the low-stringency data set yielded the lowest average gene tree–species tree conflict of 0.563 among the other data sets. In our simulations, the highest average gene tree–species tree conflict observed was 0.507, by setting theta = 0.1 and alignment length = 300 bp. Therefore the lowest empirical gene tree–species tree discordance was still significantly higher than the simulated conditions with extremely high level of ILS and gene tree estimation error (*p*-value = 2.2e-16, Student’s *t*-test, Fig. 4a). The same conclusion also applies when simulating gene trees directly from species tree without setting theta *a priori* (Fig. 4b).

**Figure 4.**
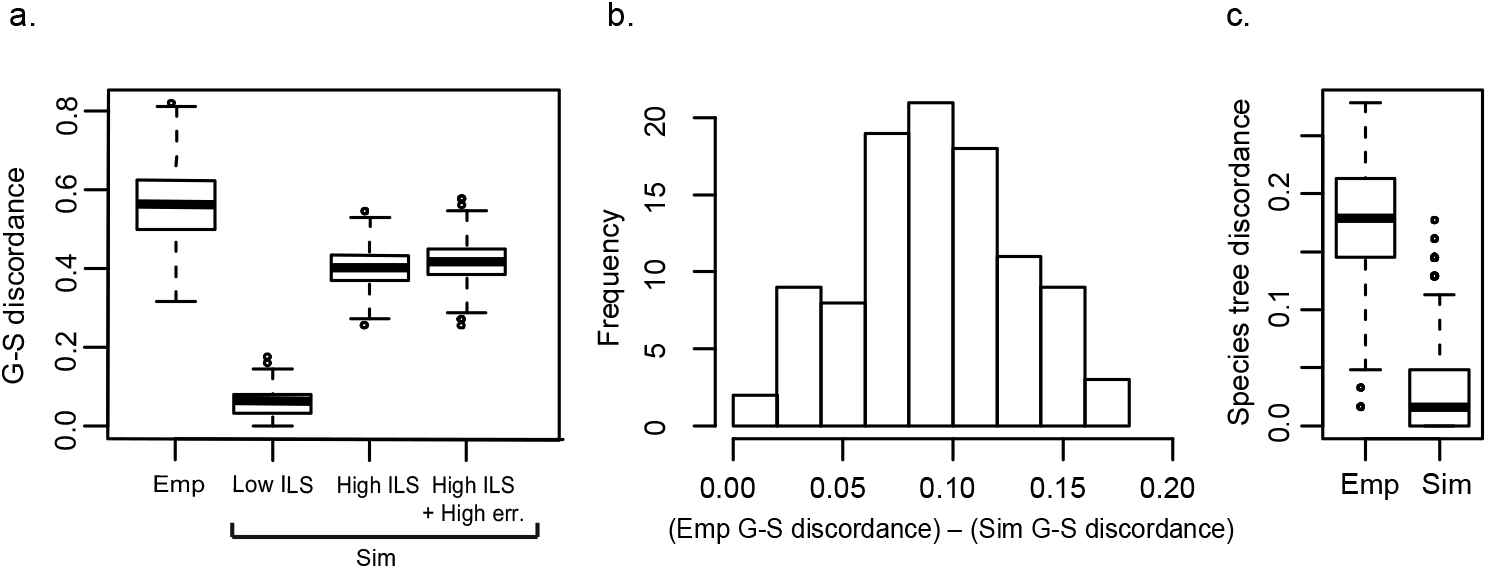
Extensive gene tree discordance in empirical versus simulated data. (a) Gene tree–species tree (G-S) discordance in the empirical (Emp) and simulated (Sim) data assuming fixed theta in simulation. Discordance is measured by RF distance between inferred gene trees and the species tree. Under various simulated conditions of ILS (e.g., ‘Low ILS’, theta = 0.01 and ‘High ILS’, theta = 0.1) and gene tree estimation error (‘High ILS + High err.’, theta = 0.1, alignment length=300bp), the simulated gene tree–species tree discordance is significantly lower than that from empirical data. (b) Gene tree–species tree discordance is higher in empirical versus simulated conditions without setting theta a priori. For each BP data set, gene tree–species tree discordance is measured and compared in both empirical and simulated data sets. Positive values indicate higher gene tree–species tree discordance in our empirical data. (c) Species tree estimation discordance in empirical data (left) and simulated data (right).

Moreover, even under such simulated conditions of extremely high ILS and gene tree estimation error, both concatenation and coalescent-based methods yielded consistent and accurate species tree estimation with no more than 12 nodes (< 0.10 RF distance, Fig. 4c, Fig. S5) failing to be recovered. The performance of coalescent-based methods is mainly affected by gene tree estimation error (Fig. S5). Under the highest gene tree estimation error (300bp), both ASTRAL and MP-EST require 1000 loci to recover the true species tree. For concatenation methods, ML estimations are robust under low ILS levels, which is consistent with previous findings (Mirarab et al. 2014b; Tonini et al. 2015). We were able to recover the correct species tree with the smallest data set (100 loci with 300bp in length) under low ILS (theta = 0.01). However, major challenges and inaccurate species trees are generated under high ILS. Under such conditions, it requires the largest data set (1500 loci with ≥400 bp length) to recover the true species tree (Fig. S5).

### MSC Model Fitness and the Relative Contribution of ILS, Gene Tree Estimation Error, and Gene Flow to Gene Tree Variation

Among all 41,664 triplets we examined, 553 (1.3%) have significant asymmetrical minor frequencies. The node with the highest Reticulation Index is the MRCA of Clade 1 and Clade 2 (the MRCA of Salicaceae and Euphorbiaceae; Fig. 5c). 10.3% of the triplets associated with this node are significantly asymmetrical. According to our relative importance decomposition analysis, ILS, gene tree estimation error, and gene flow explain 57.5% of the gene trees variation using the lmg algorithm (R^2^ = 0.575). When scaling these three factors to sum 100%, gene tree estimation error is the most dominant factor, which explains 52% of the gene tree variation (Fig. S6). The second most significant factor is gene flow, which explains 32% of the gene tree variation. And ILS explains the least variation (15%). The relative ranks of these three factors are consistent among regression methods and bootstrap replicates (Fig. S6).

**Figure 5.**
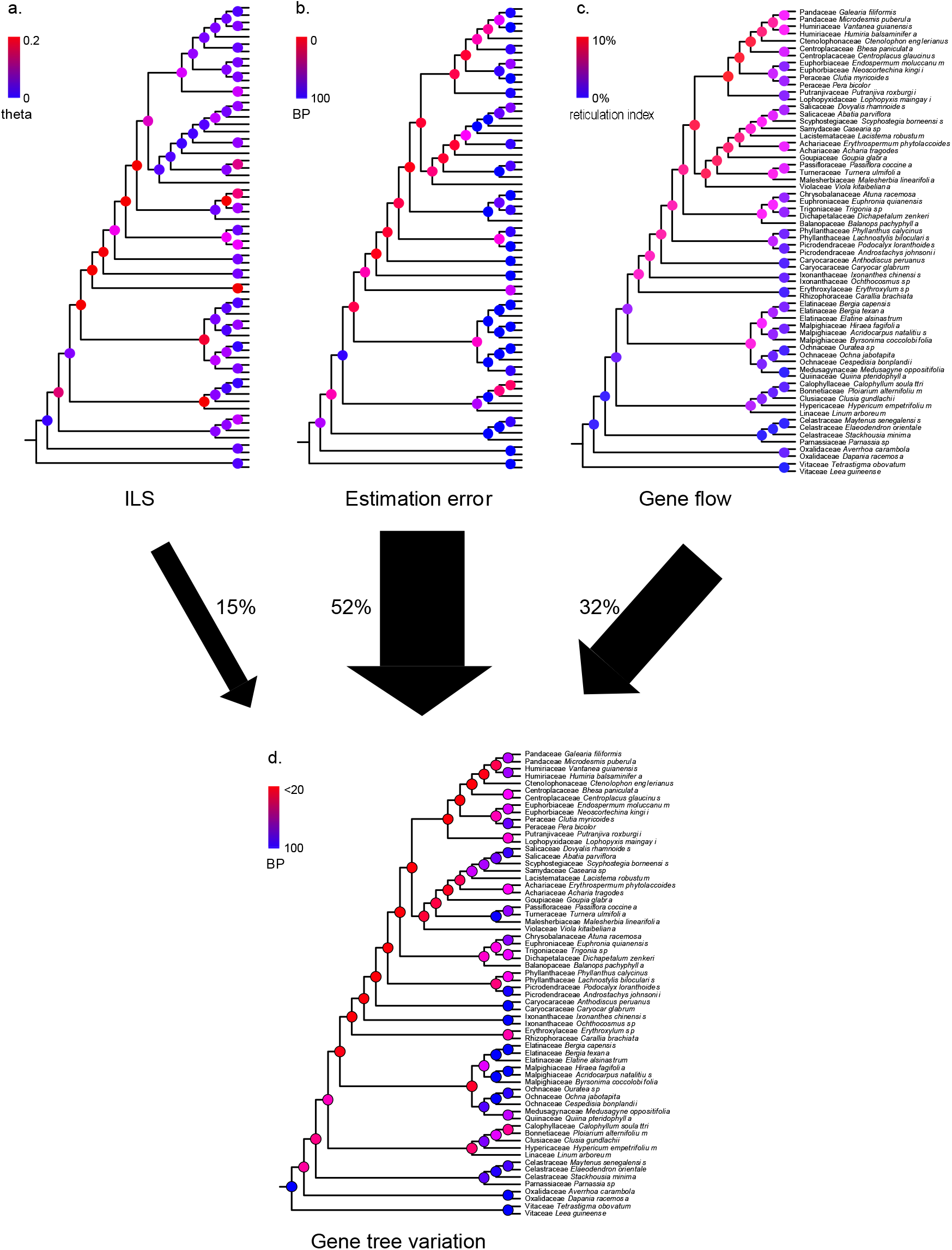
Relative contributions of ILS, estimation error, and gene flow across Malpighiales. (a) ILS. Nodes are colored by inferred population mutation parameter theta. (b) Gene tree estimation error. Nodes are colored by BP values, which represent percentage of recovered nodes from simulation (see Materials and Methods). (c) Gene flow. Nodes are colored by Reticulation Index. (d) Gene tree variation. Nodal BPs reflect nodal recovery in gene trees. Percentages of gene tree variation ascribed to ILS, estimation error, and gene flow are indicated by black arrows.

Further investigation revealed significant negative correlation (*p*-value 2.2e-16) between the overall gene tree variation and species tree resolution (Fig. S7a). All of the contributors to gene tree variation—ILS, tree estimation error, and introgression—are strongly negatively correlated with species tree resolution (*p*-value <6.6e-4). We observed the highest level of ILS, introgression and gene tree estimation error for the most recalcitrant nodes along the backbone of the phylogeny using our methods (Fig. S7b–d). This further corroborates our conclusion that a combination of all three factors contribute to this low resolution. We did not find significant correlation between the estimated level of introgression and ILS, suggesting that our triplet method can effectively distinguish these two phenomena. However, both ILS and introgression are positively correlated to gene tree estimation error (*p*-value < 6.8e-3).

### The triplet-frequency-based method identified three hotspots of introgression in yeasts

Our species tree of yeast inferred using MP-EST is identical to the topology reported in the original study by Salichos and Rokas (2013). We identified 116 asymmetrical triplets among the 1,771 triplets in the yeast species tree. These triplets revealed three hotspots of introgression that correspond to those identified by Yu and Nakhleh (2015): in the MRCA of *Saccharomyces kluyveri* and *Kluyveromyces waltii*, the MRCA of *Zygosaccharomyces rouxii* and *Saccharomyces castellii*, and the MRCA of *Candida guilliermondii* and *Debaryomyces hansenii* (Fig. S8). The first two hotspots of reticulation (the MRCA of *S. kluyveri* and *K. waltii*, the MRCA of *Z. rouxii* and *S. castellii*) reflect the donor and recipient lineage of one of the two reticulation branches identified by Yu and Nakhleh (2015). The third introgression hotspot involving the MRCA of *C. guilliermondii* and *D. hansenii* reflects the second reticulation branch inferred in Yu and Nakhleh (2015).

## Discussion

Our results indicate that despite extensive phylogenomic data, the early branching order of Malpighiales remains uncertain. We attribute this to a combination of factors—a perfect storm—involving ILS, gene tree estimation error, and gene flow. Below we highlight our findings in four subsections: the phylogenetic utility of flanking regions in sequence capture data, novel phylogenetic relationships gleaned for Malpighiales, an efficient method to investigate gene flow in large data sets, and a novel simulation-based method to decompose gene tree variation into various contributing factors.

### Flanking Regions Greatly Enhance Phylogenetic Resolution

Hybrid enrichment probes are designed to capture highly conserved anchor regions as well as the more variable flanking regions adjacent to these anchors. Despite the perceived utility of these flanking regions in mammals (McCormack et al. 2012) and more recently in in plants (Fragoso-Martínez et al. 2017), assumptions of the enhanced phylogenetic utility of these flanking regions have not been tested explicitly to our knowledge. Here, we observed significantly higher average ML BP across gene trees, increased species tree resolution, and most importantly, increased species tree estimation congruency as flanking regions were increasingly added (Fig. S2). This suggests that longer loci, favoring more phylogenetically informative flanking regions, should be prioritized in future anchored hybrid enrichment kit designs. These flanking regions represent genomic regions under nearly neutral selection where mutation rates are high, and thus appear to be a rich source of phylogenetic utility. It has been demonstrated that the inclusion of genes with higher mutation rates can greatly enhance phylogenetic resolution, even deep within organismal phylogenies (Hilu et al. 2003; Lanier et al. 2014). Our site-subsampling strategy, which includes increasingly larger proportions of these more rapidly evolving flanking regions provides the first empirical evidence that these regions are particularly informative for resolving phylogenetic relationships at shallow and deeper phylogenetic depths.

### Sequence Capture Data Confirms Malpighiales Relationships and Identifies Novel Clades

We assessed the performance of hybrid enrichment markers by evaluating support for major clades previously identified from plastome sequences (Xi et al., 2012; Fig. 3). The majority of the well-supported (>90 BP) clades identified by Xi et al. (2012) are corroborated in our analyses with high confidence (>97 BP). These include the parietal, clusioid, phyllanthoid, ochnoid, chrysobalanoid, and putranjivoid subclades. With rare exception, relationships within these clades were also identical to those by Xi et al. (2012). In the case of the parietal and clusioid clades, internal resolutions were less well supported owing to conflicting topologies recovered among coalescent and concatenation methods (low nodal support indicated by ‘–’ in Fig. 3). Within the parietal clade, for example, the monophyly of the salicoids *sensu* Xi et al. (2012, Fig. 3) is supported by the RAxML phylogeny with moderate support (69 BP) but is not supported in any of the coalescent methods.

Additionally, we discovered several noteworthy clades that conflict with those reported by Xi et al. (2012). The euphorbioids, malpighioids, and rhizophoroids were paraphyletic in all of the best resolved MP-EST, ASTRAL, and RAxML analyses (Fig. 3). The euphorbioids—including Euphorbiaceae, Peraceae, Lophopyxidaceae, Linaceae, and Ixonanthaceae—were split into four polyphyletic groups. In particular, Linaceae was placed as sister to the clusioid clade in all of the best resolved coalescent and concatenation analyses (Fig. 3). The affiliation of Linaceae to the clusioids instead of to other members of the euphorbioids is also supported in a recent transcriptomic study of this group with less dense taxon sampling (Cai et al. 2019). Within malpighioids, Centroplacaceae is confidently placed (>86 BP) with Humiriaceae, Pandaceae, and Ctenolophonaceae (Fig. 3) instead of with Malpighiaceae and Elatinaceae. This relationship is partially supported by Wurdack et al. (2004) in which Centroplacaceae was placed with Pandaceae, although with low support. Within the rhizophoroids, Ctenolophonaceae was well nested (>98 BP for coalescent methods) within a clade including Euphorbiaceae and Pandaceae (Clade1 in Fig. 3) rather than with Rhizophoraceae and Erythroxylaceae.

### ILS and Gene Tree Estimation Error Alone Are Insufficient to Explain the Lack of Species Tree Resolution in Malpighiales

Our simulations to explore gene tree heterogeneity encompass the full distributional range of ILS and gene tree estimation error inferred from the empirical data, and clearly demonstrate that the data we have assembled should be sufficient to resolve Malpighiales species tree relationships. Specifically, despite our inability to estimate a well-resolved species tree from our empirical data, we were able to recover a species tree with very high confidence in simulation (mean nodal support >91 BP). This is true even when ILS (theta = 0.1) and gene tree estimation error (alignment length = 300bp) were set to the highest levels inferred from our empirical data. Such extreme levels of theta, in particular, are ten times higher than empirical estimations from *Arabidopsis* and *Drosophila* (0.01– 0.001 in both cases; Drost and Lee 1995; Fischer et al. 2017). Even when down sampling our data set under these extreme conditions to include a mere 100 loci, both concatenation and coalescent analyses recover the true species with no more than 10% error (Fig. S5). In addition, we observed far fewer conflicts among species trees reconstructed from different methods and data partitions in simulation versus from those estimated from the empirical data (Fig. 4c). These results suggest that ILS and gene tree estimation error alone are insufficient to explain the lack of resolution along the spine of Malpighiales, and suggest that additional factors likely contribute to gene tree heterogeneity.

### Gene Flow Compromises Malpighiales Species Tree Resolution: A Novel Method for Assessing Gene Tree Heterogeneity

Beyond ILS and gene tree estimation error, gene tree heterogeneity is also attributable to two other common biological factors: gene duplication and gene flow (Yang 2006). As we demonstrate above, the first two factors alone are insufficient to explain this lack of resolution. Orthology assignment problems owing to gene duplications are also highly unlikely for two reasons. First, our sequence capture data set was specifically designed for single copy nuclear loci across land plants (Buddenhagen et al. 2016). Second, large-scale genome duplication identified in Malpighiales all occurred *subsequent* to the explosive radiation where discordance is localized (Cai et al. 2019). Thus, biased gene loss arising from genome duplications are unlikely to hinder our ability to resolve backbone relationships in the order. Additional analytical artifacts not reflected in our assessment include homolog calls, alignment error, and most importantly, misspecification of DNA substitution models, all of which can compromise species tree estimation. Though these analytical errors may explain some discordance, it is quite possible that conflicts are attributed to additional biological phenomena.

Gene flow has yet to receive attention in phylogenomic studies, especially at deep-time phylogenetic scales. It is estimated that at least 25% of plant species and 10% of animal species hybridize (Mallet 2007) and various network inference methods have been developed to assess gene flow in phylogenies (Nakhleh 2013). These methods have provided valuable insights into reticulate evolution, including those associated with the rapid radiations in wild tomatoes and heliconius butterflies (Pease et al. 2016; Edelman et al. 2019). However, the performance of these methods often relies on accurate species tree estimation and the generation of a handful of alternative species tree topologies to conduct hypothesis testing. However, when alternative topologies are too numerous to evaluate, such as along the backbone of Malpighiales, existing tools become quite limited. In particular, these methods are computationally expensive and amenable only to small data sets. For example, maximum likelihood can only be applied to networks involving fewer than 10 taxa and three reticulations (Yu and Nakhleh 2015). We leveraged the theoretical predictions of triplet frequencies to make inferences about gene flow by summarizing the distribution of lineages involved in horizontal processes using our novel measurement statistic, the Reticulation Index. Our method can effectively identify hotspots of reticulate evolution, including both the donor and recipient lineage, in large clades and in deep time, and provide valuable guidance to empirical studies. We further validated the application of our Reticulation Index using the yeast data set from Salichos and Rokas (2013). The three hotspots we identified in the yeast phylogeny (Fig. S8) correspond precisely to the two reticulation branches previously inferred by Yu and Nakhleh (2015), thus demonstrating the promise of our method for applications in larger phylogenies like Malpighiales.

In Malpighiales, the Reticulation Indices are especially high in deeper parts of the phylogeny, suggesting that certain clades may contribute substantially to this phenomenon (Fig. 5c). In particular, we hypothesize that the overabundance of asymmetrical triplets observed within Clades 1 (MRCA of Euphorbiaceae and Putranjivaceae) and Clade 2 (MRCA of Salicaceae and Violaceae) result from ancient and persistent gene flow between early diverging members of these lineages. Specifically, Clade 1 contains six paralogous lineages from the plastid phylogeny (Xi et al. 2012) and is a major hotspot for plastid-nuclear conflict. Such conflict is widely recognized as an indicator of introgression (Soltis and Kuzoff 1995; Baum et al. 1998). Moreover, members of two clades, the putranjivoids and Pandaceae, have previously been implicated in the top three most unstable nodes of all angiosperms (Smith et al. 2013). We hypothesize that this may be attributed to the chimeric nature of their ancestral genealogy resulting from gene flow. The Reticulation Index is also significantly negatively correlated with species tree resolution (Fig. S7d), suggesting that introgression is an important barrier for robust species tree estimation in Malpighiales. For the most recalcitrant nodes where almost no bootstrap replicates recover the same topology, we also observed the highest values of inferred introgression. In the meantime, no correlation is identified between the estimated level of ILS and introgression, suggesting that our methods can effectively distinguish ILS and introgression. However, nodes with strong introgression signals also have higher gene tree estimation error (Fig. S7e). One possible explanation for such correlation is that the short branch lengths created by introgression may lead to elevated estimation error at these nodes.

To better characterize gene tree variation attributable to ILS, gene tree estimation error and gene flow, we devised a novel regression method to parse variation attributable to these analytical and biological factors. Our method of decomposing gene tree variation revealed that the majority of variation is due to estimation error (52%), while gene flow and ILS account for 32% and 15%, respectively. This decomposition analysis is based on estimations of ILS, gene tree error, and gene flow through simulation and is subject to common limitations of regression analyses. As a result, errors from the simulation and regression analysis can render the absolute values of these percentages less reliable. Regardless, the relative influence of biological and analytical aspects of gene tree variation as interpreted from these metrics can shed important light on empirical investigations and the development of enhanced species tree inference methods. For example, though gene tree error is to blame for the majority of gene tree variation in our test case, gene flow still plays a significant role in gene tree variation. Therefore, a species network inference method that accommodates gene flow is essential to better understand the early evolutionary history of Malpighiales. Application of this method to other taxonomic groups will also reveal the key factors contributing to recalcitrant relationships and provide guidance for phylogenomic marker design targeting at specific questions.

Our results suggest that a confluence of factors—ILS, gene tree estimation error, and gene flow—influence this lack of resolution and contribute to a perfect storm inhibiting our ability to reconstruct branching order along the back of the Malpighiales phylogeny. Gene flow, in particular, is a potentially potent, and overlooked factor accounting for this phenomenon. Despite a relatively small percentage of asymmetrical triplets attributed to gene flow (1.3% of all triplets), they appear to contribute substantially to gene tree heterogeneity based on our relative importance decomposition (32%). Our approach of interrogating this phenomenon using triplet frequencies and the relative importance analyses can elucidate factors that give rise to gene tree variation. These approaches are likely to be especially useful for investigating the causes of recalcitrant relationships, especially at deeper phylogenetic nodes, and to highlight instances where relationships are better modeled as a network rather than a bifurcating tree.

## Acknowledgements

We would like to thank Sean Holland and Michelle Kortyna at the Florida State University Center for Anchored Phylogenomics for their assistance with data collection and analysis. We would like to thank Scott Edwards and Davis Lab members for helpful comments. Funding for this study came from Harvard University, and US National Science Foundation Assembling the Tree of Life Grant DEB-0622764, and from DEB-1120243, and DEB-1355064 (to C.C.D.).

## Figure Captions

**Figure S1** Number of PI sites and mean gene tree BP is positively correlated with alignment length in high/medium/low-stringency data sets. (a,b) Correlation between number of PI sites (a) or mean gene tree BP (b) with alignment lengths inferred from the high-stringency data set. (c,d) Correlation between number of PI sites (c) or mean gene tree BP (d) with alignment lengths inferred from the medium-stringency data set. (e,f) Correlation between number of PI sites (e) or mean gene tree BP (f) with alignment lengths inferred from low-stringency data set. Pearson’s R^2^ is presented at lower right corner of each plot.

**Figure S2** Increased gene tree and species tree resolution as more flanking sites are included in the analysis. (a) Distribution of mean gene tree BP in high/medium/low-stringency data sets. (b,c) Increased species tree BP in concatenation (b) and coalescent analysis (c). Analyses with same locus subsampling are connected by lines. (d) Increased species tree inference consistency reflected by pairwise RF distance.

**Figure S3** Species tree discordance is more sensitive to site and locus subsampling in coalescent (black) versus concatenation analyses (grey). Left, distribution of pairwise species tree distances derived from all coalescent (black) and concatenation analyses (grey) measured by RF distance. Right, distribution of pairwise species tree distances from coalescent (black) and concatenation (grey) analyses measured by weighted RF (WRF) distance (weighted by nodal support).

**Figure S4** Gene tree estimation error in empirical and simulated data. Gene tree estimation error is measured by RF distance to the ‘true gene tree’ for both empirical and simulated data sets. In both cases, gene tree estimation error is negatively correlated with alignment length.

**Figure S5** Species tree estimation error in simulated data sets. Species tree estimation error is measured by RF distance from inferred species in each analysis to the known species tree. Results derived from alignments of varying lengths (300, 400, 500, 1000, 1500 bp) are marked by different color and shape. (a,b) Species tree estimation error of ExaML under low (a) and high (b) ILS. (c,d) Species tree estimation error of MP-EST under low (c) and high (d) ILS. (e,f) Species tree estimation error of ASTRAL-II under low (e) and high (f) ILS.

**Figure S6** Relative importance of ILS, gene tree estimation error, and gene flow in generating gene tree variation based on four regression methods. Percentages are normalized to sum 100%. 95% confidence intervals are represented by bars.

**Figure S7** ILS, gene tree estimation error, and introgression contribute to low species tree resolution in Malpighiales. Species tree resolution is represented by nodal support from the MP-EST phylogeny in Figure 3 from the main text. The other statistics reflect the variables presented in Figure 5. The *p*-value of the Pearson’s correlation test is indicated in the upper right corner in each panel. (a) Significant negative correlation between gene tree variation and species tree resolution. (b) Significant negative correlation between ILS and species tree resolution. (c) Significant negative correlation between gene tree estimation error and species tree resolution. (d) Significant negative correlation between introgression and species tree resolution. (e) Significant positive correlation between gene tree estimation error and introgression. (f) No significant correlation between gene tree estimation error and ILS.

**Figure S8** Hotspots of reticulate evolution in baker’s yeast. Species phylogeny is inferred from MP-EST with the 1,070 genes trees in Salichos and Rokas (2013). Nodes are colored by Reticulation Index. Black thick arrows indicate inferred reticulation by Yu and Nakhleh (2015) for comparative purpose.

**Table S1** Voucher and GenBank information for 64 species in Malpighiales, Celastrales, Oxalidales, and Vitales used for anchored hybrid enrichment.

**Table S2** Species tree estimation strategies using various phylogenetic estimation methods and phylogenetic subsampling methods (see Supplementary Note 1) with high-, medium-, and low-stringency data sets.

**Table S3** Coalescent and mutational parameters of simulated data sets.

**Table S4** Summary statistics of 423 loci in high/medium/low-stringency data sets, including number of captured taxa, alignment length, number of PI sites, and mean gene tree BP.

