## Supplementary figures and images for "The Perfect Storm: Gene Tree Estimation Error, Incomplete Lineage Sorting, and Ancient Gene Flow Explain the Most Recalcitrant Ancient Angiosperm Clade, Malpighiales"

### Fig. S1

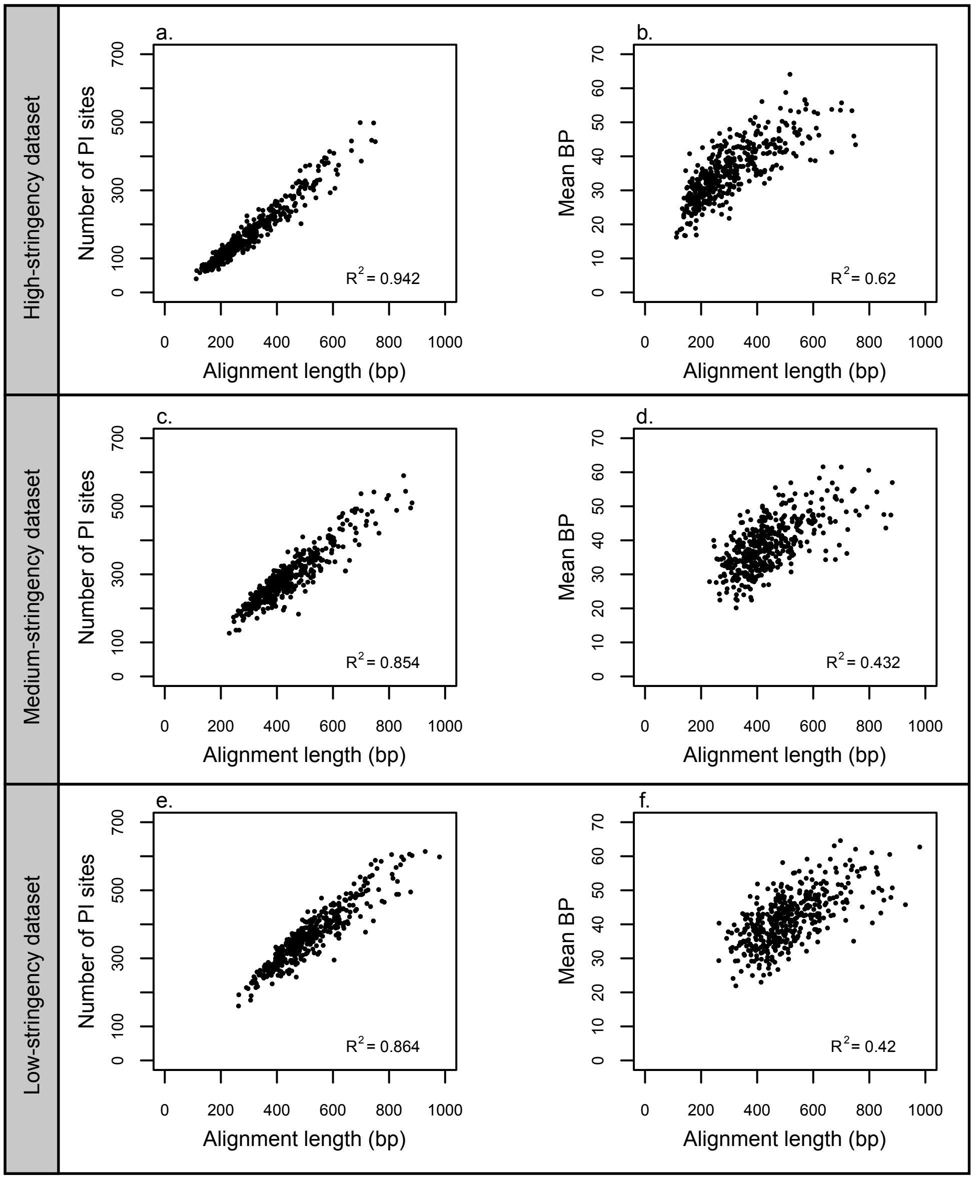

### Fig. S2

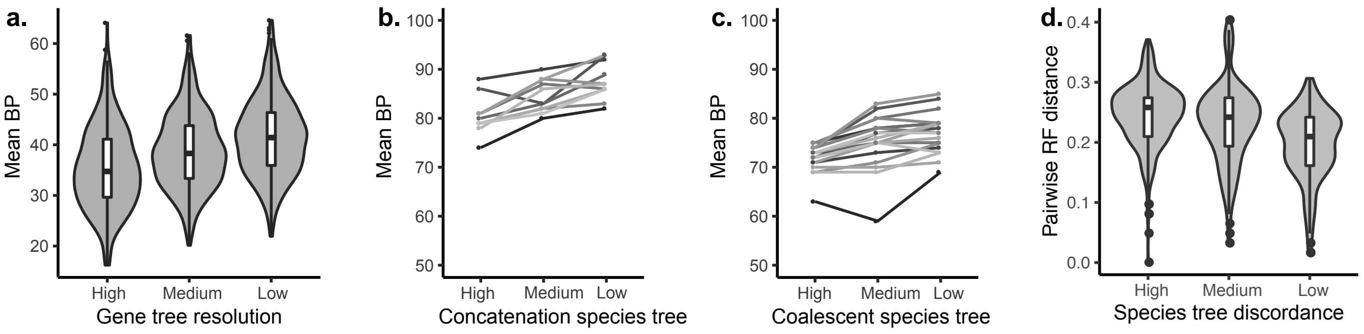

### Fig. S3

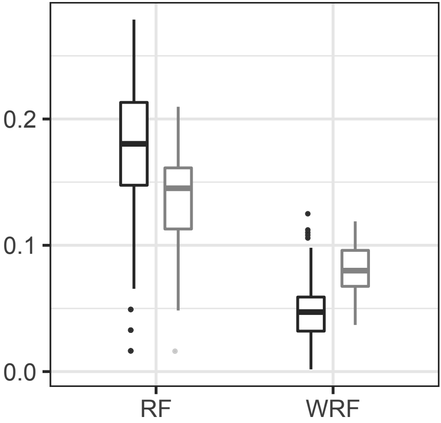

### Fig. S4

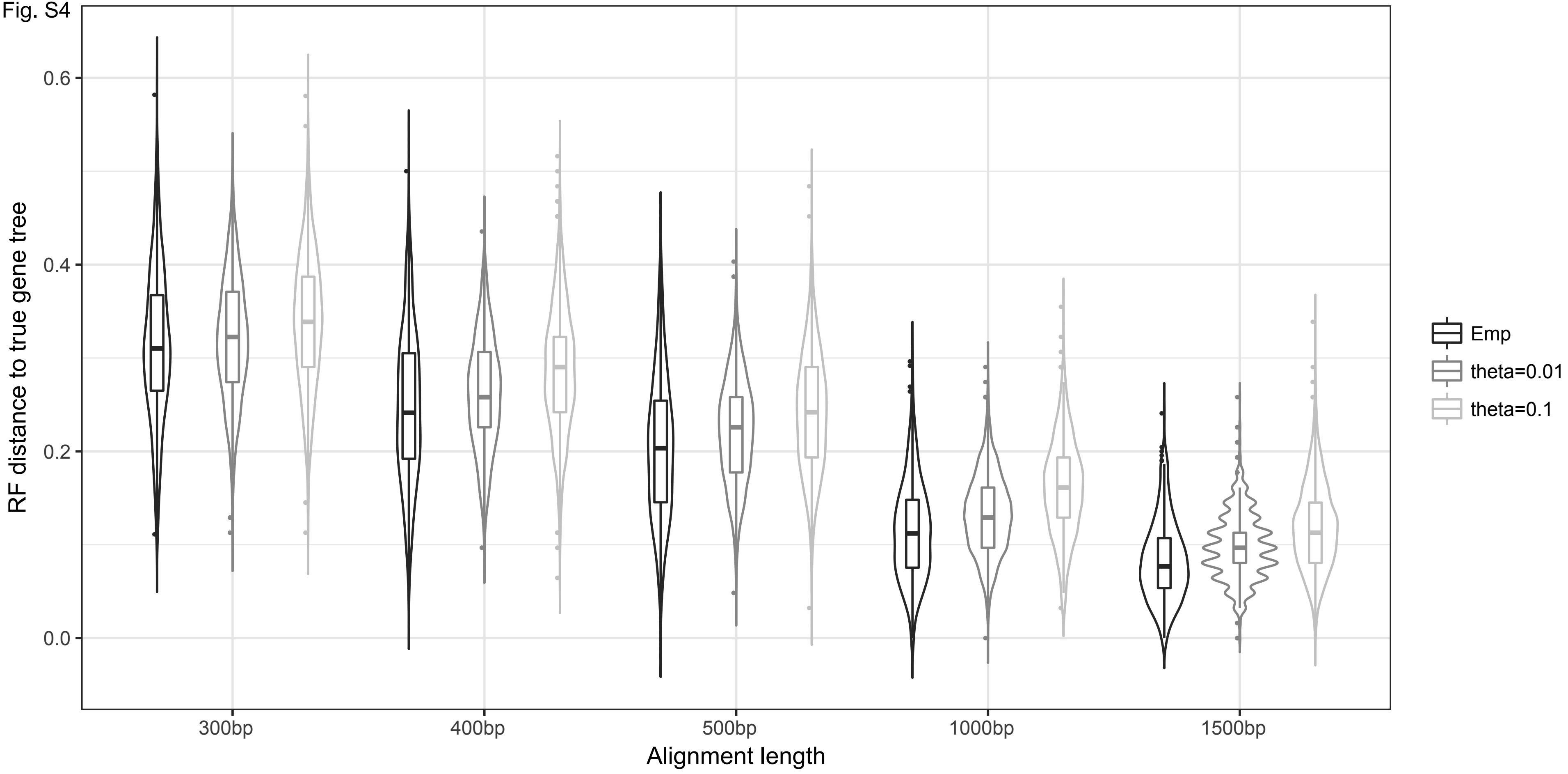

### Fig. S5

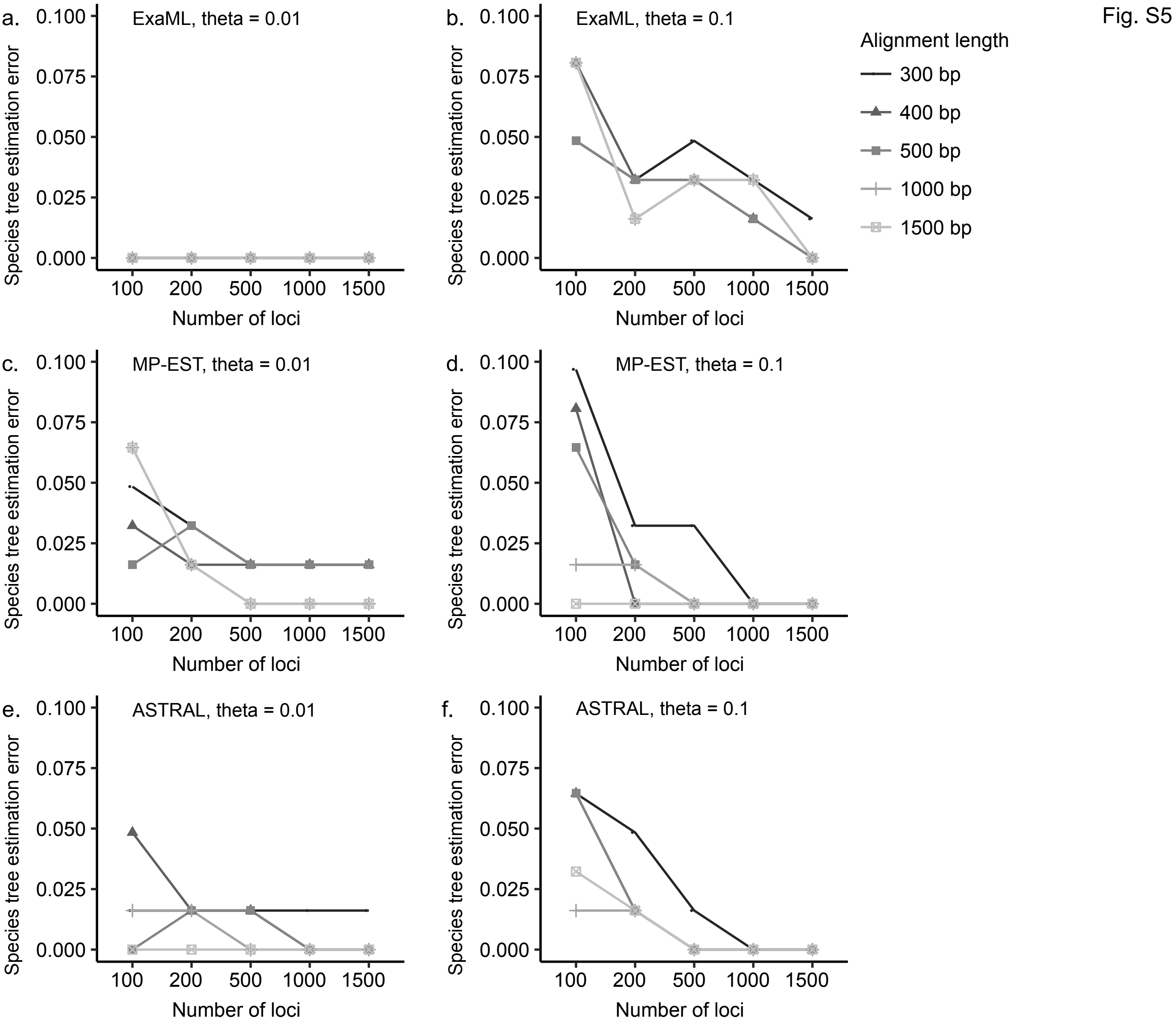

### Fig. S6

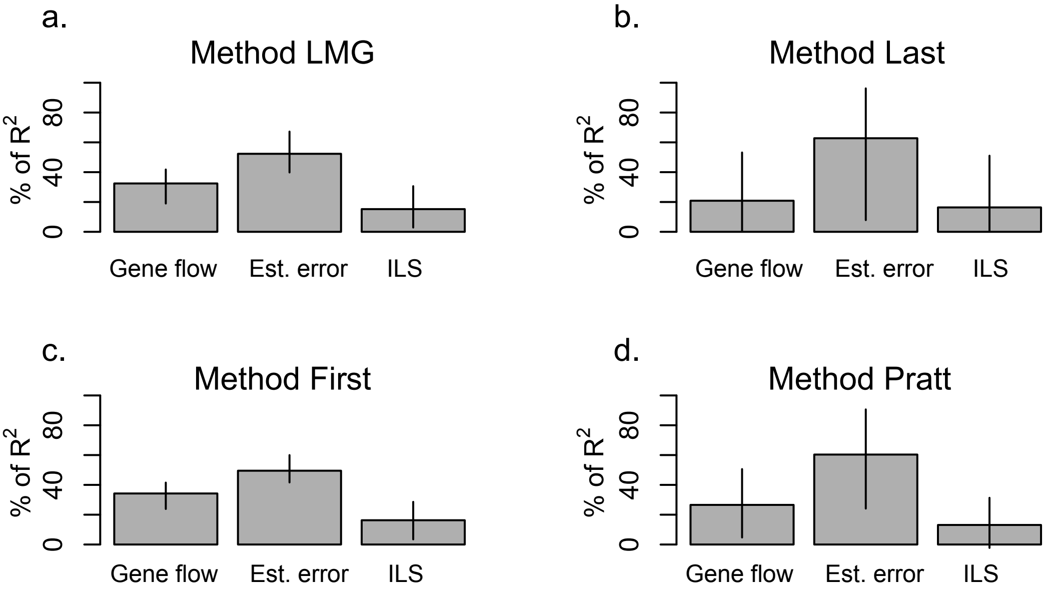

### Fig. S7

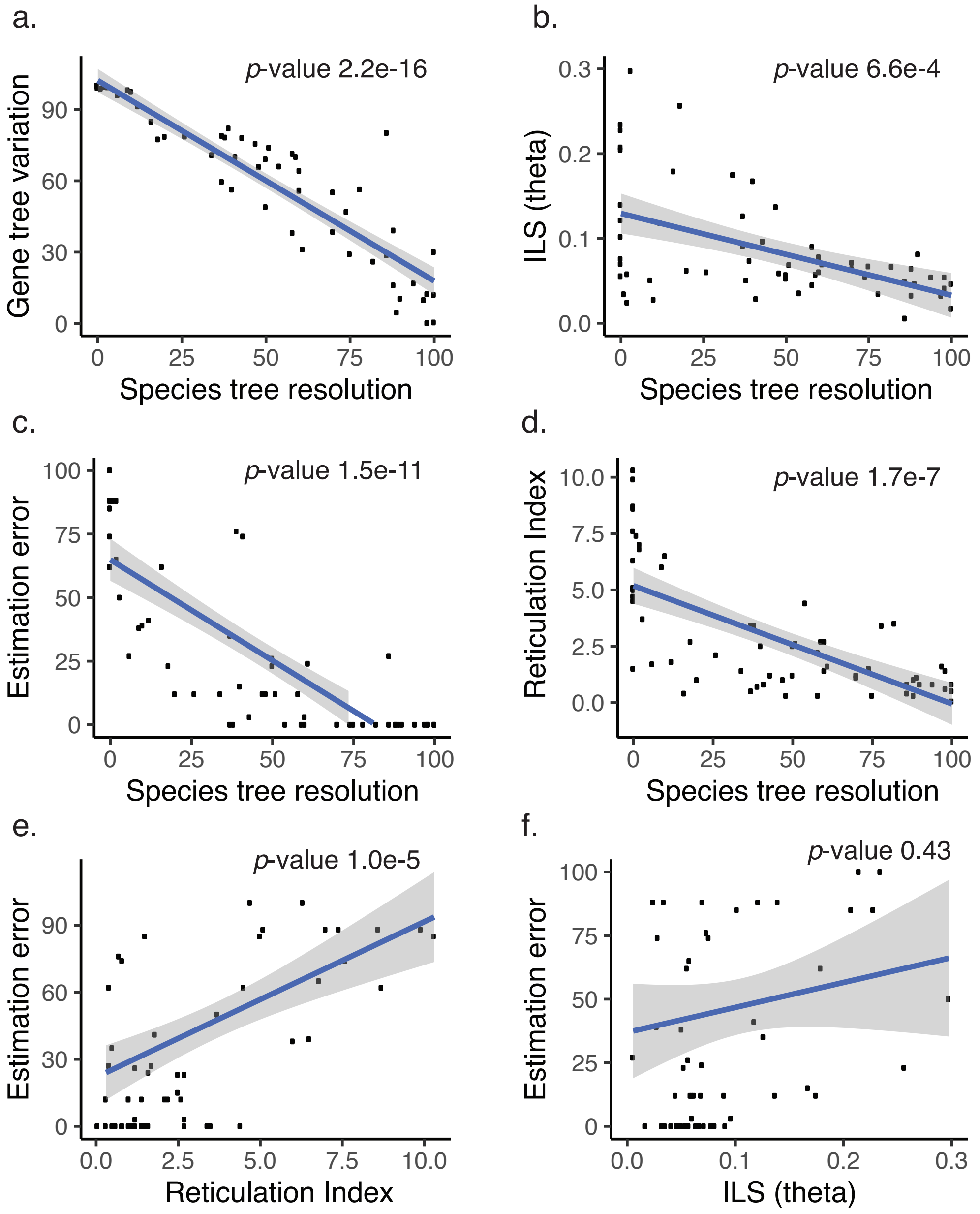

### Fig. S8

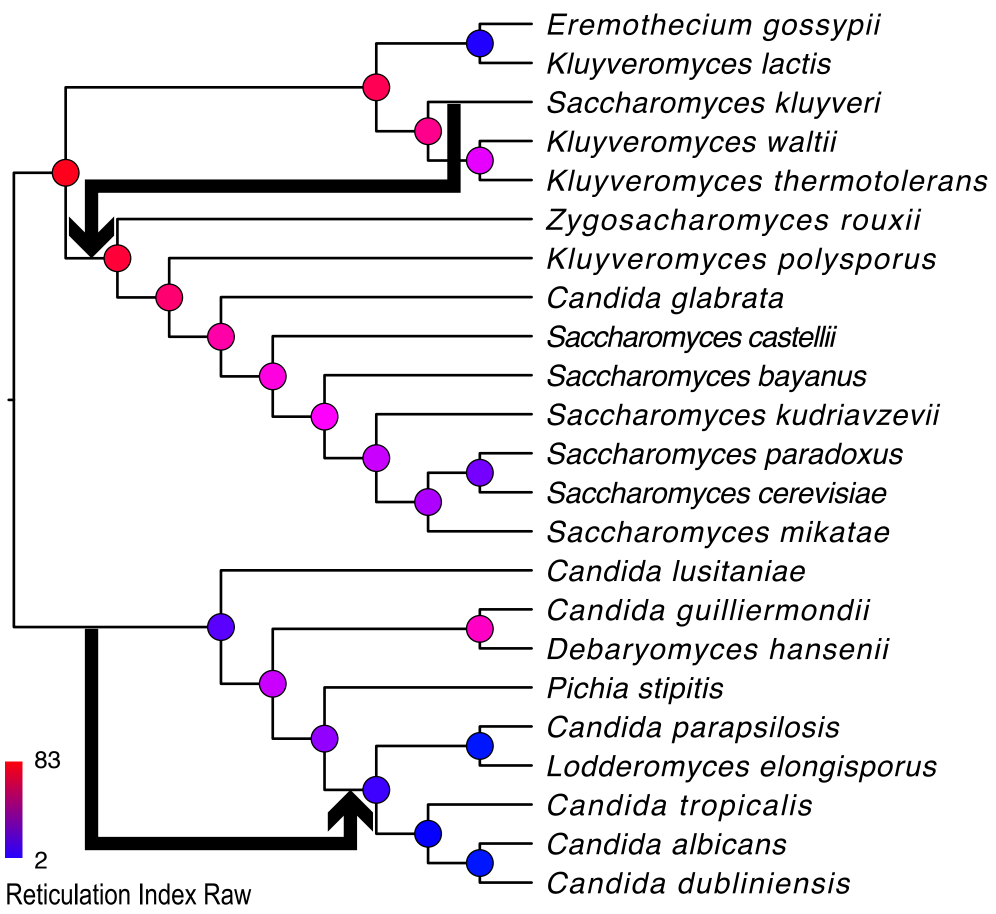
